# Expected 10-anonymity of HyperLogLog sketches for federated queries of clinical data repositories

**DOI:** 10.1101/2021.01.30.428918

**Authors:** Ziye Tao, Griffin M. Weber, Yun William Yu

## Abstract

**Motivation:** The rapid growth in of electronic medical records provide immense potential to researchers, but are often silo-ed at separate hospitals. As a result, federated networks have arisen, which allow simultaneously querying medical databases at a group of connected institutions. The most basic such query is the aggregate count—e.g. How many patients have diabetes? However, depending on the protocol used to estimate that total, there is always a trade-off in the accuracy of the estimate against the risk of leaking confidential data. Prior work has shown that it is possible to empirically control that trade-off by using the HyperLogLog (HLL) probabilistic sketch.

**Results:** In this article, we prove complementary theoretical bounds on the k-anonymity privacy risk of using HLL sketches, as well as exhibit code to efficiently compute those bounds.

**Availability:** https://github.com/tzyRachel/K-anonymity-Expectation

**Supplementary information:** N/A

## 1 Introduction

Clinical data containing patients’ personal medical records are important resources for biomedical research. Fully centralizing that data may permit the widest array of potential analyses, this is often not feasible due to privacy and confidentiality requirements (Heatherly *et al*., 2013; Benitez and Malin, 2010; Emam *et al*., 2009). During times of pressing need, such as during a global pandemic, these privacy requirements may be justifiably relaxed Haendel *et al*. (2020)—such as using trusted 3rd party vendors such as Datavant (Kho and Goel, 2019)—but even then, it is important to keep in mind the various privacy-utility trade-offs (Bengio *et al*., 2021, 2020). A more privacy friendly alternative is to use a federated network instead, which give hospitals control over their local databases; then, a distributed query tool enables researchers to send queries to the network, such as “how many patients across the network have diabetes” (Weber, 2015; Brat *et al*., 2020). A number of these hospital networks have emerged, including the Shared Health Research Information Network (SHRINE) for Harvard affiliated hospitals (Weber *et al*., 2009), the Federated Aggregate Cohort Estimator (FACE) developed through a collaboration of five universities and institutions (Wyatt *et al*., 2014), the open-source PopMedNet (Davies *et al*., 2016), and the Patient Centered Outcomes Research Institute (PCORI) launched PCORnet as a network of networks (Fleurence *et al*., 2014).

However, patients often receive medical care from multiple hospitals, so medical records at different hospitals may be duplicated or incomplete. Depending on the aggregation method used to combine results from the network, this can produce errors. For example, consider using a simple summation of aggregate counts: if a patient with hypertension receives medical care from both Hospital A and Hospital B, then it is possible that the sum will double count that patient, which results in the overestimation of the number of patients with hypertension (Weber, 2013).

Of course, this problem can be mostly alleviated by sending a hashed identifier of patients matching each hospital’s queries to a trusted third party, but that again raises privacy concerns (Oechslin, 2003). There is some natural trade-off between the privacy guaranteed to individual patients and the accuracy of the aggregate query, and hashed identifiers and simple summation fall at opposite ends of the spectrum. Several of the authors of this paper recently proposed using the HyperLogLog ‘probabilistic sketch’ Durand and Flajolet (2003); Flajolet and Martin (1985); Flajolet *et al*. (2007) to access intermediate trade-offs of privacy vs. accuracy (Yu and Weber, 2020) Probabilistic counting was introduced to the computer literature decades ago, and has found use in analyzing large streaming data in a variety of settings, ranging from internet routers (Cai *et al*., 2005) to text corpora comparisons (Broder, 1997) to genomic sequences (Ondov *et al*., 2016; Baker and Langmead, 2019; Solomon and Kingsford, 2018). Instead of sharing a single aggregate count, or sharing the full list of matching patient IDs (Weber, 2013), each hospital instead shares a smaller ‘summary sketch’ built from taking the logarithm of a coordinated random sample of *m* matching patient hashed IDs Yu and Weber (2020). Because only m patient IDs are used, and those are obfuscated through taking a logarithm, these HyperLogLog sketches are significantly more private than sending a full list of matching IDs. Due to the way the estimators work, HyperLogLog sketches have an error of about 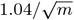, which can be much less than expected from simple summation.

But when any data is shared by a hospital to a 3rd party, no matter the method, there is risk of accidental leakage. For example, consider the case where a hospital finds that there is only one patient satisfying the criterion for a query. If this hospital returns the aggregate count as one, then this unique patient’s personal information is linked and can potentially be re-identified through a linkage attack (Yu and Weber, 2020; Emam and Dankar, 2008). To properly compare the privacy of various methods of data aggregation, we turn to the concept of k-anonymity. The basic idea behind k-anonymity is that if a method or dataset is k-anonymous, then each patient is similar to at least *k* – 1 other patients with respect to potentially identifying variables, so that it is hard to determine the identity a single patient in the data set (Emam and Dankar, 2008; Sweeney, 2002). Although other mathematical formalisms like differential privacy (Dwork, 2008) are much stronger, they are harder to work with, as they require injecting deliberate noise, and are not currently widely in use by clinical databases.

In this paper, we will assume that hospitals in a federated network implement the HyperLogLog(HLL) algorithm for queries. We will then prove bounds on the expected k-anonymity of the shared sketches, as well as provide fast algorithms for computing that expected k-anonymity. This study is an extension of previous work (Yu and Weber, 2020), which operated under the same setting and assumptions, but only provided empirical results and no proofs on the levels of privacy achieved. Here, we provide rigorous theoretical justification for those empirical claims.

## 2 Methods

### 2.1 Setting and summary

In this paper, we adopt the HyperLogLog (HLL) sketch federated clinical network setting given in prior work (Yu and Weber, 2020). For completeness, we duplicate the salient points below.

Assume that every patient has a single invariant ID that is used across hospitals. Prototypically, one might consider using social security numbers in the USA for that purpose. Even without a single unique identifier, it is possible to generate an ID based off a combination of other possibly non-unique IDs, such as first and last name, zip code, address, birthdate, etc. Unfortunately, there may be errors in these records due to character recognition errors(e.g., S and 8), phonetic errors (e.g. ph and f), and typographic errors including insertion, transposition and substitutions. Luckily, there is a lot of existing literature on this problem, and methods such as BIN-DET and BIN-PROB (Durham *et al*., 2010) have been proposed to deal with the issue. Thus, in this paper, we will treat this problem as out-of-scope and assume for simplicity that every patient has a unique stable ID known to all institutions.

Further assume that there is a federated network of hospitals (or other institutions) responding to clinical queries, along with a central party that manages and relays messages. When hospitals receive a query, they generate a list of the IDs of patients who match the query. Each hospital will use a publicly known hash function to first pseudorandomly partition the patients into m buckets and then assign a uniform pseudorandom number between 0 and 1 to each patient. The hospital then stores the order of magnitude of the smallest number within each bucket, and sends these m smallest bucket values to the central party. By applying the HLL estimator, the central party is then able to compute the aggregate count for the query with a relative error of around 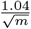 (Flajolet *et al*., 2007).

Here, we focus on an individual hospital, and want to determine the expected exposure to accidentally disclosing private information if the central party is compromised. As the HLL sketch aggregates information within each of the *m* buckets, our goal is to compute the expected number of buckets which are not k-anonymized. In line with common practice, we set *k* = 10 for most of our results, though the algorithms and proofs hold for other k. Below, we provide two approximation formulas for the expected value and in the Results section construct a table for the user to determine which approximation should be chosen based on the number of distinct patients and other relevant parameters.

### 2.2 k-anonymity and HyperLogLog

#### 2.2.1 High-level overview

The HyperLogLog (HLL) (Flajolet *et al*., 2007) probabilistic sketching algorithm is widely used to estimate the cardinality (number of different elements) of a set. Assume we have a database of electronic medical records; we can estimate the number of distinct patients by applying the HLL algorithm. The basic idea behind HLL is that the minimum value of a collection of random numbers between 0 and 1 is inversely proportional to the size of the collection. Therefore, we can estimate the cardinality of a set by first applying a hash function which maps all the elements uniformly onto [0, 1] and considering the minimum value. For the purposes of this paper, we will operate in the random oracle model, where we assume that the hash function actually maps to a random number; in practice, a standard hash function like SHA-256 would probably be employed. In order to increase the accuracy of estimation, we randomly divide the set into m partitions and then estimate the cardinality of the original set by the harmonic mean from m partitions. Furthermore, the HLL algorithm only needs to store the position of the first 1 bit in the 64-bit hash value, rather than the full patient ID hash, providing partial privacy protection. As the expected error in the final estimate is around 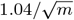, increasing m can reduce the error of HLL query but increases the risk of privacy leaks.

In our setting, when a hospital is sent a query, there are two relevant sets to consider: (1) the background population (often, the set of all patients at the hospital), and (2) the set of patients matching the query. The reason for considering the background population is that they can ‘hide’ patients who match the query by providing plausible deniability. The hospital will return a HLL sketch, which contains *m* values—the maximum position of the first 1 bit within each bucket. We define a HLL bucket with value *x* to be ‘k-anonymous’ if at least *k* – 1 patients in the background population have hash value *x*; we will call these corresponding hash values in the background population hash *collisions* (Yu and Weber, 2020). This means that if an attacker gets access to the sketch and can narrow down the set of potential patients to the background population, they cannot determine with certainty which of the *k* patients with that hash value was in the set of patients matching the query. Our goal is to determine the expected number of buckets that are not at least 10-anonymous (Figure 1).

**Fig. 1:**
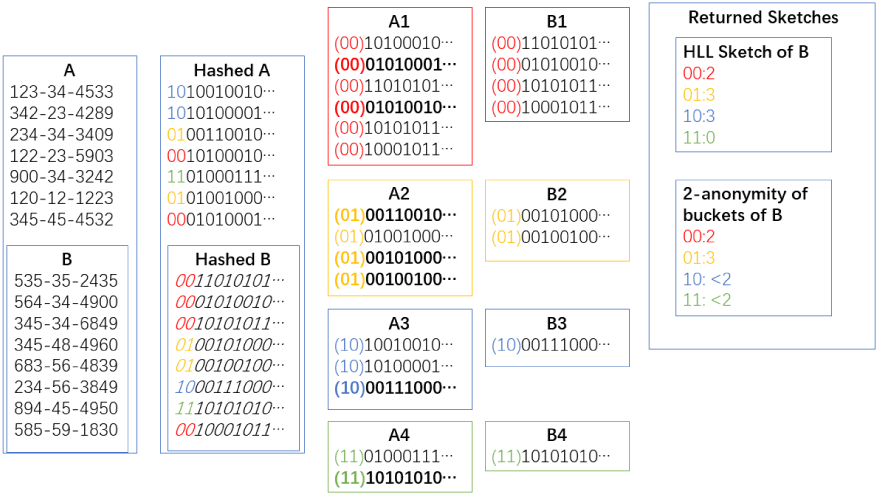
Illustration of HyperLogLog k-anonymity. A hospital has an identified set *B* contained within the background population *A*. Binary hashes are taken of all patient identifiers. Those hashes are first used to partition the patients into 4 buckets. Within each bucket of *B*, the smallest value is chosen as the representative. Then the k-anonymity of that bucket is the number of hashes in the corresponding bucket of the background population that share the same position of the leading 1 bit.

#### 2.2.2 Formal description

Let’s recast the textual description above a bit more rigorously as the following mathematical problem:

Let *A* be a set and *B* ⊆ *A* is a non-empty subset of *A. A* represents the background population and *B* represents patients satisfying the query. We define 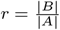 as the ratio of number of patients satisfying the query to background population (also sometimes known as concept prevalence).

Let *σ*: *A* → (0, 1] be a one-way hash function. In theory, we assume that we have a shared oracle available to both parties. In practice, a cryptographic hash function such as SHA-1, SHA-224, or SHA-256 (Johnson, 2020) is generally used. *σ* uniformly maps each element in A to a random real number in the interval (0,1].

Let 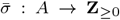 be defined by 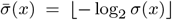. This function returns the number of 0 bits before the first 1 bit in *x* ∈ (0,1] under a base-2 expansion.

Let *p*: *A* → {1,…, *m*} be a map that randomly partitions patients into *m* buckets. In practice, this map can also be derived from a cryptographic hash function. From the partition function *p*, we define *A_i_* = {*x* ∈ *A* | *p*(*x*) = *i*} and *B_i_* = *B* ⋂ *A_i_*, which respectively represent the *i*th bucket in whole database and sample.

Let 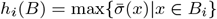 be the maximum number of zeros before the first one among all hash values represented in base-2 in the ith bucket of B which is *B_i_*.

Let 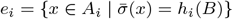 be the set of elements in the *i*th bucket of A which collide with the elements in *B_i_*.

We want to compute the 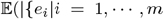 and 0 < |*e_i_*| ≤ *k* – 1}|), the expected number of non k-anonymous buckets.

### 2.3 Probability of <k-anonymity without partition function

As described above, we need to consider the collisions against all m buckets. Here, however, we first show a simple analysis with no partition function (i.e. the case where *m* = 1) and compute the probability of each possible number of collisions so that in the later sections we can use this result to compute the desired expected value of ‘non k-anonymous’ buckets.

Since there is only one bucket, there are only two sets A and B which represent the set of all patients and the set of patients matching the query respectively. We denote 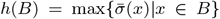, the maximum number of zeros before the first one among all hash values in base-2 in *B*, and 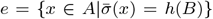, the set of collisions. We want to compute the probability that the number of collisions is less or equal to *k*, which is *P*(|*e*| ≤ *k* | *A, B*).

Each element in *σ*(*A*) can be thought of as an i.i.d. random variable with distribution *Unif* (0,1). Therefore, 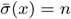 if and only if 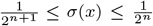. Then we get 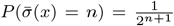. Thus, 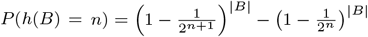.

#### Lemma 2.1.

*Given sets B* ⊂ *A, the probability of exactly* 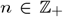 *collisions is:*

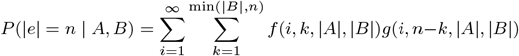

*where* 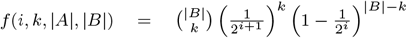 *and* 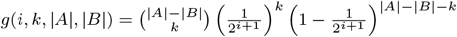.

Proof. Since the sets A and B are fixed, we use *P*(|*e*| = *n*) to represent *P*(|*e*| = *n* | *A,B*) for notational simplicity here.

By the law of total probability, we know that 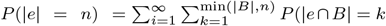 and *h*(*B*) = *i*) *P*(|*e* ⋂ (*A* – *B*) | = *n* – *k* and *h*(*B*) = *i*)

First we consider the case where we have *k* collisions in *e* ⋂ *B*:

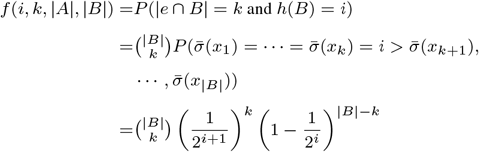

Next we consider the case where we have *k* collisions in *e* ⋂ (*A* – *B*):

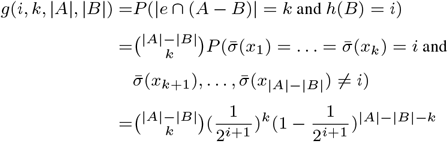

Thus

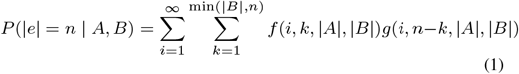

and 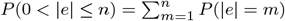

### 2.4 Expected number of buckets with less than *k* collisions

Recall that *A* is the background population and *B* the set of patients satisfying the query criteria. We denote the buckets of *A* and *B* under our partition function by *A*_1_, ···, *A_m_* and *B*_1_, ···, *B_m_* where *B_i_* = *B* ⋂ *A_i_* for *i* = 1,.., *m* and *e_i_* is the sets of collisions in the ith bucket. Thus, the expected value of the number of buckets with no more than *k* collision is *E*(|{*e_i_* | |*e_i_*| ≤ *k, i* = 1, ···, *m*}|).

Note that (|*A*_1_|, ···, |*A_m_*|) ~ *Multinomial*(|*A*|, *p*_1_, ···, *p_m_*) with 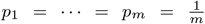. Therefore, we know for a single bucket, say *A*_1_, its cardinality follows a binomial distribution that is 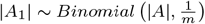

With a given *A_i_*, |*B_i_*| ~ *Hypergeometric*(|*A*|, |*A_i_*|, |*B*|). Thus,

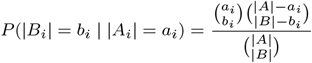

where *b_i_* ∈ {0, 1, ···, min(*a_i_*, |*B*|)} and 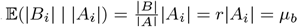 and 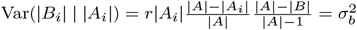.

#### Theorem 2.2.

*The expected number of buckets which have at least 1 collision but no more than k collisions is:*

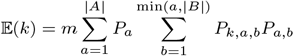

*where P_a_* = *P*(|*A*_1_| = *a*), *P_a,b_* = *P* (|*B*_1_| = *b* | |*A*_1_| = *a*) *and P_k,a,b_* = *P* (0 < |*e*_1_| ≤ *k* | |*A*_1_| = *a*, |*B*_1_| = *b*).

Proof.

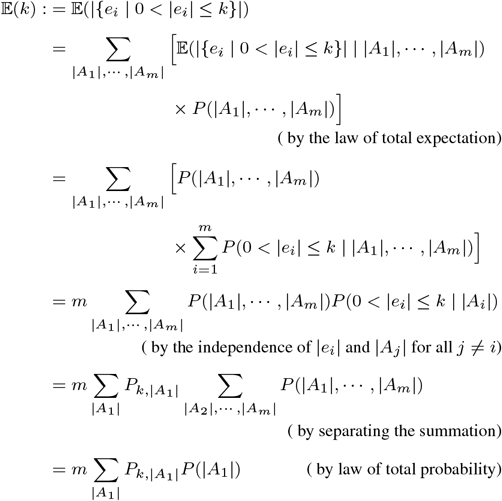

where *P*_*k*,|*A*_1_|_ = *P*(0 < |*e*_1_| ≤ *k* | |*A*_1_|).

In order to compute *P*(0 < |*e*_1_| ≤ *k* | |*A*_1_|), we have to consider the range of |*B_i_*| which is {0, 1, ···, min(|*B*|, |*A*_1_)}.

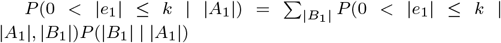.

In contrast to the simple case in section 2.3, here *B*_1_ is not necessarily a proper subset of *A*_1_ because *A*_1_ can be the empty set and thus *B*_1_ is also an empty set in this case. The collision number is zero if and only if *A*_1_ is an empty sets. Therefore, we will expand the formula in lemma 2.1 to compute *P*(0 < |*e*_1_| ≤ *k* | |*A*_1_|, |*B*_1_|). Furthermore, if we want rule out the case of zero collisions—because when the bucket is empty, there is not a patient ID for which we need to guarantee k-anonymity—we should set the range of |*A*_1_| and |*B*_1_| as {1, 2, ···, |*A*|} and {1, 2, ···, min(*a*, |*B*|)} respectively.

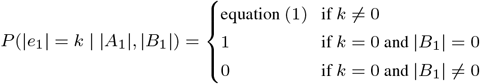

Therefore, we will get:

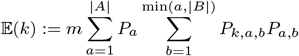

where 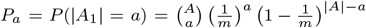, 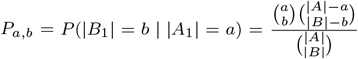 and *P_k,a,b_* = *P*(|*e*_1_| ≤ *k*| |*A*_1_| = *a*, |*B*_1_| = *b*).

## 3 Algorithms

### 3.1 Time Complexity of evaluating expectation

Again, recall that *A* is the background population, *B* is the set of patients satisfying the query criteria, and e is the set of collisions. In section 2.4, we gave an explicit formula for computing *P*(|*e*| ≤ *k* | |*A*|, |*B*|). However, the time complexity of carrying out that computation is troublesome.

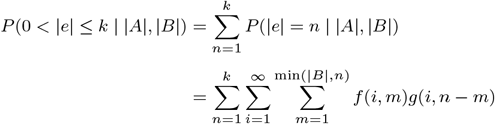

where 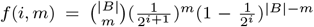 and 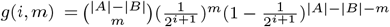

Usually, *k* is smaller than |*B*| and the infinity in the second sum will be replaced by 64 (or some other constant <100) because it represents the maximum number of zeros before the first one among all hash values in base 2. As there are only 7 billion people on Earth, 64-bits is sufficient for the original hash function to have low probability of collisions. Therefore, the time complexity is 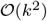 for at most k collisions.

We consider the time complexity of computing the desired expectation. Theoretically, the range of |*A*_1_| is {1, 2, ···, |*A*|} and the range of |*B*_1_| is {1, 2, ···, min(|*B*|, |*A*_1_|)}. Therefore, the computation time is:

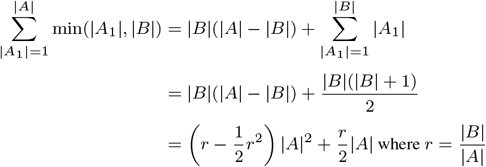

and the time complexity is 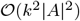 which is quadratic in the size of population for at most k collisions. In practice, for large set sizes, it is computationally infeasible to use this theoretical formula to compute the desired expectation; thus, in the remainder of this paper we analyze fast approximations.

### 3.2 Approximation A1: Concentration inequalities

When |*A*| is large, it is impossible to sum over whole range of |*A*_1_|. Therefore, we will use concentration inequalities to restrict |*A*_1_| and |*B*_1_ | to a smaller range. Because there is only an exponentially small probability that *A*_1_ and *B*_1_ will fall outside these restricted windows, this will have minimal effect on the final answer while reducing the computation time from quadratic to linear in the size of *A*.

Recall that 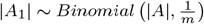 and 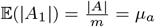, 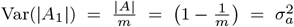. In order to reduce the time complexity, we will restrict |*A*_1_| in our computations to the interval (*L_a_, U_a_*):= (*μ_a_* – 5*σ_a_, μ_a_* + 5*σ_a_*).

Recall that |*B*_1_| ~ Hypergeometric(|*A*|, |*A*_1_|, |*B*|) for a given |*A*_1_| and 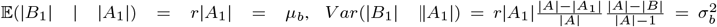. However, we define 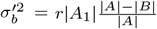 which is greater than 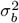 and restrict |*B*_1_| in the interval 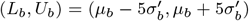 in order to compute the error bound more easily below in Section 3.2.1. After concentration, we can make sure that 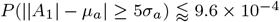 and 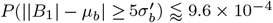 which is shown below in detail in Section 3.2.1. As an aside, while these two intervals of |*A*_1_| and |*B*_1_| have been chosen for analysing the error bound and time complexity analytically, in the computing code we can directly use built-in functions to compute the relevant confidence intervals for |*A*_1_| and |*B*_1_|.

By the concentration inequalities on |*A*_1_| and |*B*_1_|, the desired expectation will be approximated by:

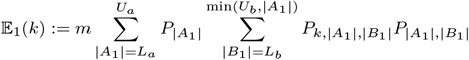

where *P*_|*A*_1_|, |*B*_1_|_ = *P*(|*B*_1_| | |*A*_1_|), *P*_|*A*_1_|_ = *P*(|*A*_1_|) and *P*_*k*, |*A*_1_|, |*B*_1_|_ = *P*(0 < |*e*_1_| ≤ *k* ||*A*_1_|, |*B*_1_|)

The computation time after concentration is:

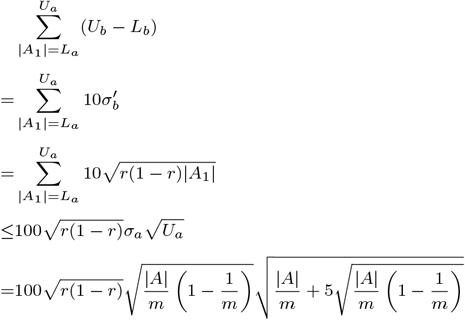

So, the time complexity after concentration is 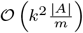 which is linear in 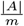. After concentration, the expected value *E*_1_(*k*) is smaller than the actual 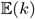, but we can bound the error.

#### 3.2.1 Error bounds

Recall that 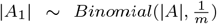 and 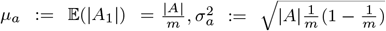. We concentrate |*A*_1_| in the interval (*L_a_, U_a_*):= (*μ_a_* – 5*σ_a_, μ_a_* + 5*σ_a_*). We define *F_a_*(*x*):= *P*(|*A*_1_| ≤ *x*) the cumulative density function of |*A*_1_|.

First we consider the concentration on |*A*_1_|. We will apply the higher moments inequality on |*A*_1_| — *μ_a_* (Blum *et al*., 2020):

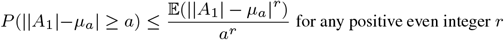

If we choose *r* = 6 then, we will get:

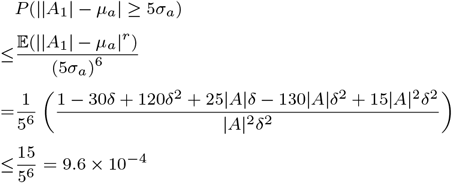

where 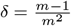

Then we consider the |*B*_1_| for a given |*A*_1_|. For a given |*A*_1_|, we know |*B*_1_| ~ *Hypergeometric* (|*A*|, |*A*_1_|, |*B*|) and 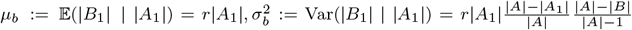. We concentrate |*B*_1_| in the interval 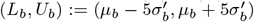 where 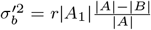. We define *F_b_*(*x*):= *P*(|*B*_1_| ≤ *x*| |*A*_1_|) the cumulative density function of |*B*_1_| for a given |*A*_1_|.

Note that for *X* ~ Binomial(|*A*_1_|, *r*), we can get 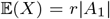 and Var(*X*) = *r*(1 – *r*)|*A*_1_|. The expected value is equal to 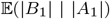 and the variance is equal to 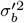 which is bigger than the variance of |*B*_1_| for this given |*A*_1_|. This explains that the hypergeometric distribution is more concentrated about the mean than the binomial distribution (Kalbfleisch, 1985). Therefore, we will use this binomial distribution to bound the tail of our hypergeometric distribution:

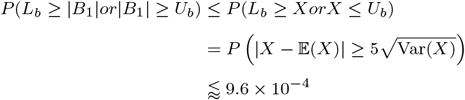

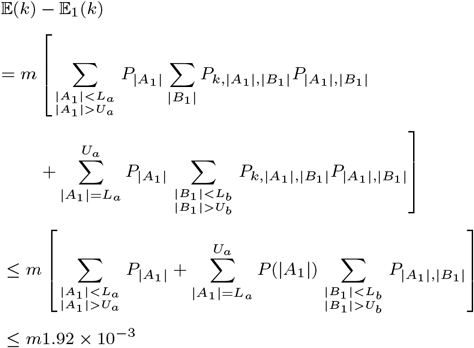

where *P*_|*A*_1_|, |*B*_1_|_ = *P*(|*B*| | |*A*_1_|), *P*_|*A*_1_|_ = *P*(|*A*_1_|) and *P*_k, |*A*_1_|, |*B*_1_|_ = *P*(|*e*_1_| ≤ *k* | |*A*_1_|, |*B*_1_|)

But the in the computing code, we can use the built-in function to find the interval (*L_a_, U_a_*) and (*L_b_, U_b_*) such that *P*(*L_a_* ≤ |*A*_1_| ≤ *U_a_*) ≥ 1 – *α* and *P*(*L_b_* ≤ |*B*_1_| ≤ *U_b_*) ≥ 1 – *α*. This will not affect the time complexity and can ensure that the absolute error between the estimated expected value and the actual expected value is < 1 by choosing a proper *α*. It is obvious the smaller *α* is, the smaller the error will be, but the intervals (*L_a_, U_a_*) and (*L_b_, U_b_*) will be bigger which means a longer computing time. Therefore, there is a trade-off between accuracy and speed (see Figure 2 for real computing time). Fortunately, in all cases we explore, the *L_a_* and *U_a_* given above can ensure that *α* < 5 × 10^-5^.

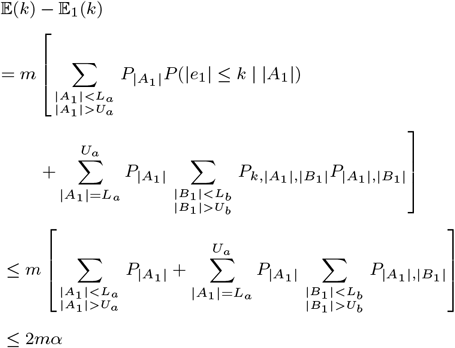

where *P*_|*A*_1_|, |*B*_1_|_ = *P*(|*B*_1_| | |*A*_1_|), *P*_|*A*_1_|_ = *P*(|*A*_1_|) and *P*_*k*, |*A*_1_|, |*B*_1_|_ = *P* (0 < |*e*_1_| ≤ *k* | |*A*_1_|, |*B*_1_|)

**Fig. 2:**
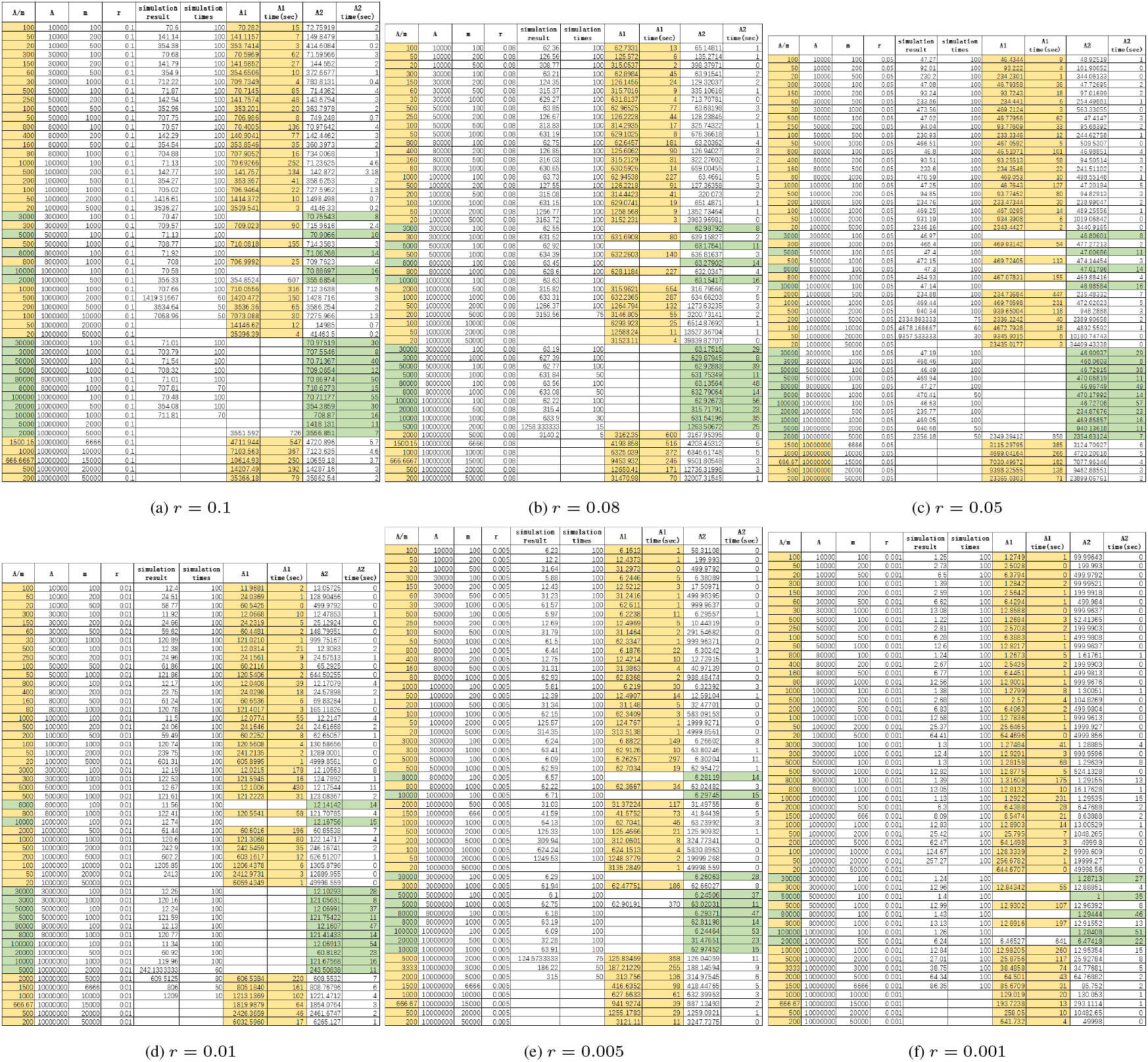
Simulation result and expected number of non 10-anonymous buckets from Approximations A1 and A2 under different combinations of m(number of buckets) when r(prevalence rate). Subfigures grouped by prevalence rate. The tables show the simulation results and computing results by the two approximations. Yellow means that we compute this case by approximation A1 and green means we compute this case by approximation A2. Full data tables are also available on Github in machine-readable format.

#### 3.2.2 Approximation A2: mean-field approximation

Although the time complexity after concentration is linear 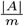, for large |*A*| and *m* small, this speedup is often still not enough. We can further approximate *P*(|*e*_1_| ≤ *k* | |*A*_1_|, |*B*_1_|) by *P*(|*e*_1_| ≤ *k* | |*A*_1_|, |*B*_1_| = *r*|*A*_1_|) and get the following approximation of the expectation:

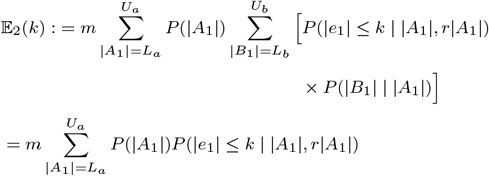

This is a ‘mean-field’ approximation based on Approximation A1. The basic idea behind this approximation is to use the probability at the mean value which is *P*(0 < |*e*_1_| < *k* | |*A*_1_|, *r*|*A*_1_|) to represent all the probabilities *P*(0 < |*e*_1_| < *k* | |*A*_1_|, |*B*_1_|) when |*B*_1_| ∈ (*L_b_, U_b_*) because *P*(0 < |*e*_1_| < *k*| |*A*_1_|, |*B*_1_|) is monotonic increasing in |*B*_1_| and the interval (*L_b_, U_b_*) is small enough compared with the theoretical range (0, min(|*A*_1_|, |*B*|)).

The range of |*A*_1_| is still (*L_a_, U_a_*) = (*μ_α_* – 5*σ_a_, μ_a_* + 5*σ_a_*). Therefore the computation time of E2 is:

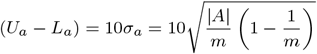

and the time complexity is 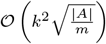. The real computing time will be discussed in the Results section. Unfortunately, we do not have a strong provable guarantee with this approximation, but it seems empirically to work well in practice.

## 4 Results

In order to assess the accuracy-speed trade-offs of our two approximations, we ran simulations measuring the empirical k-anonymity of patients in several different regimes using HyperLogLog sketches. Then, we compared those empirical values against the approximations described in this paper. In the large cardinality regimes, it is computationally infeasible to run full simulations, so we only compare the run-times of the two approximation methods. In Figure 2, we provide full tables of these results. In table 1, we provide a high-level summary giving a practitioner guidance on which method is appropriate under those particular parameter choices. All computations were run in single-thread mode on an AMD Ryzen Threadripper 3970X 32-core CPU machine running Ubuntu 18.04.5 LTS (bionic) with 256 GiB of RAM. Code is available on Github: https://github.com/tzyRachel/K-anonymity-Expectation

**Table 1.**
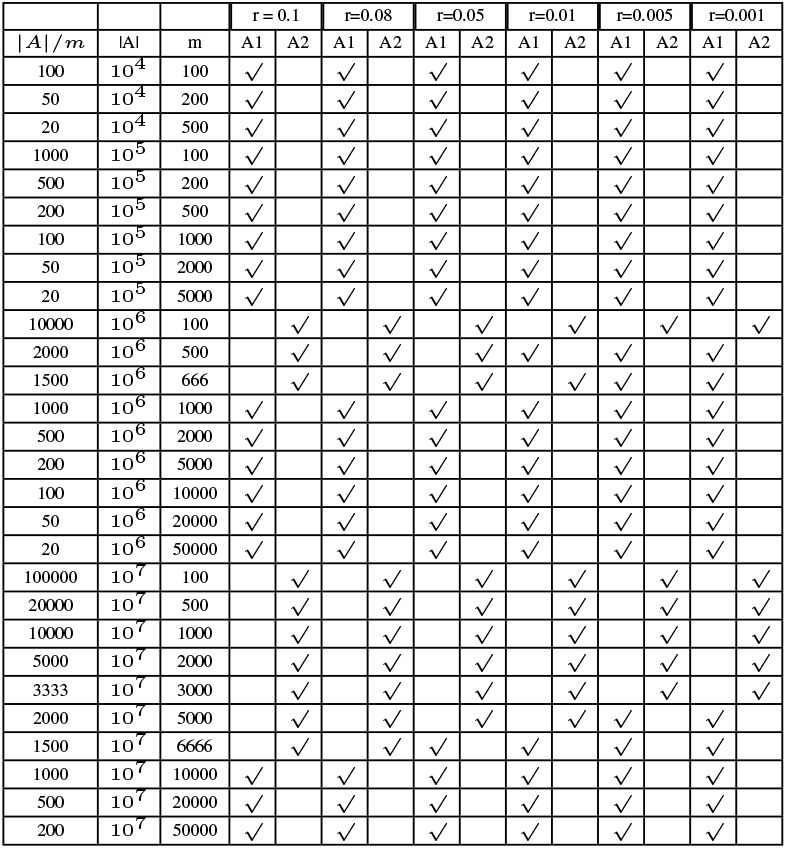
Choice Table for Approximation Method. *A* is the total size of the hospital background population, *m* is the number of buckets used in the HyperLogLog sketch, and *r* is the fraction of the background population that matches the query criteria. ‘A1’ and ‘A2’ respectively denote approximations 1 and 2. For every one of the parameter regimes, we used simulations to determine which of the approximation methods is more suitable for the practitioner.

Recall that A represents the number of all patients, *B* represents the number of patients who meet some query criteria, and *m* is the number of buckets in the HLL process. We introduce 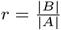 to represent the ratio of |*A*| and |*B*|, because as we will see, this ratio controls to a large extent the number of collisions. Intuitively, *r* represents the number of background population persons who could be used to provide plausible deniability to each patient in the query set.

Our simulations sweep over the different combinations of the parameters *A, r* and *m* to construct a table to fit Approximations A1 and A2. In all simulations, we restrict |*A*| in the interval [10^4^, 10^7^] and *m* in the interval [100, 50000]. Since the simulations are run under the condition of ‘10-anonymity’, we make sure that 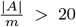 which is the mean value of the single bucket size. Also, *r* is restricted in the interval [0.001, 0.1] and we choose 6 different values of r which are: 0.1,0.08, 0.05, 0.01, 0.005, 0.001 to run the simulations and compare the simulation results with computing results.

As we discussed in the Methods section, we can estimate the desired expected value by both Approximations A1 and A2. The final choice of Approximation A1 or Approximation A2 seems to be dependent primarily on 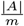. In most cases, when 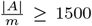, Approximation A2 is good enough and the computing time is no longer than 3 minutes. When 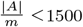, Approximation A2 will be not accurate enough and we have to choose Approximation A1. The computing time of Approximation A1 is proportional to 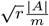, which is sometimes a concern. When 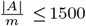, the computing time is usually no longer than 8 minutes. But there are several special cases, such as when when *r* = 0.1 and *r* = 0.08, that the computing time at *A* = 10^7^, *m* = 6666 is approximately 10 minutes which might be acceptable but is really not ideal. Furthermore, in extreme cases, the approximate expected k-anonymity return by Approximations 1 and 2 differ by about 10 (2).

To make things easier for the end-practitioner, we provide a summary ‘choice’ table (Table 1) guiding them on which approximation is suggested, based on different numbers of patients, numbers of buckets and ratios of number of patients matching query to all patients.

Figure 3 show the errors between the approximation results (based on the choice table 1) and simulation results (Figure 2) when number of distinct patients is 10^7^ and number of buckets are 100 and 1000 respectively. The absolute values of all the errors are no more than 4.

**Fig. 3:**
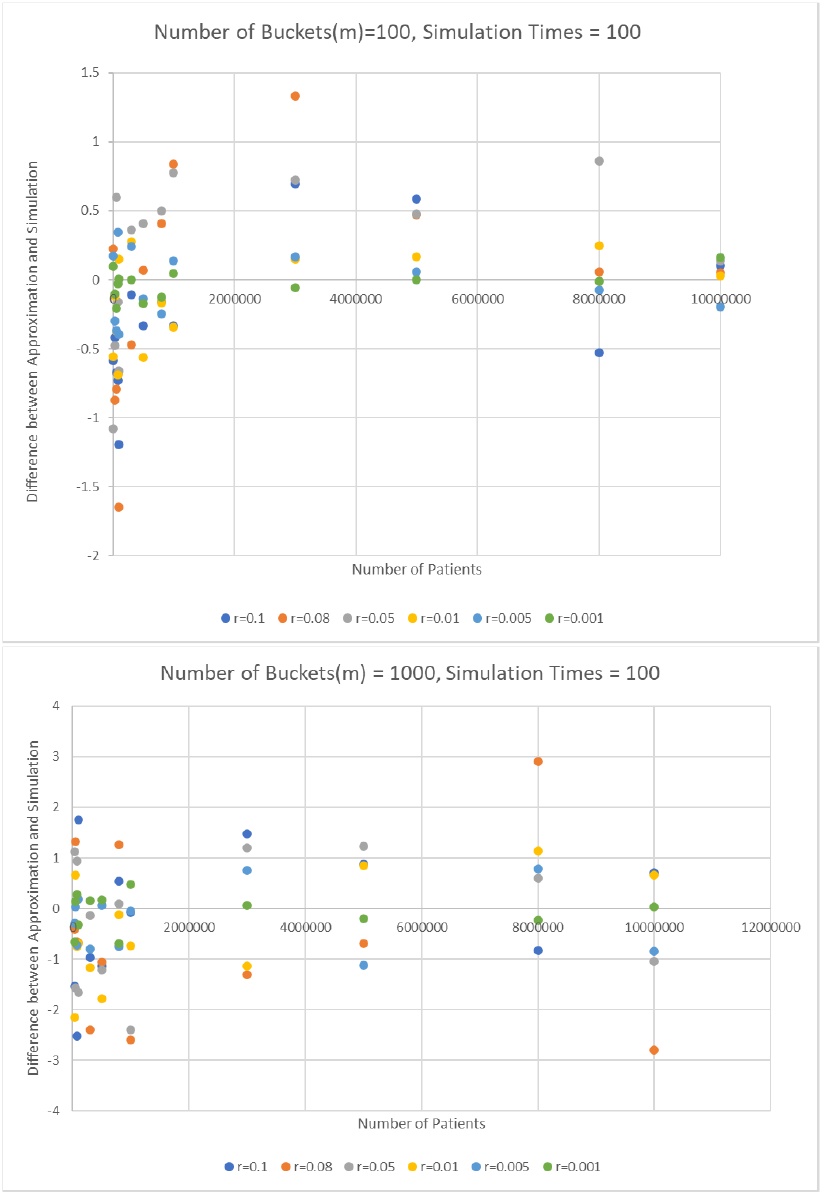
Errors between Approximation (based on choice table) and simulation of 100 random trials with number of buckets =100 (top) and 1000 (bottom).

## 5 Discussion

We first note that all of the approximations we have provided finish on the order of minutes. As they are analytical approximations, there is also no need to run them multiple times. Although we have not shown explicit simulation run-times in the tables above, the larger simulations take upwards of hours; furthermore, we did not perform simulations for the largest parameter ranges because we expected those to take significantly longer. Thus, our approximations significantly speed up the process of determining the expected privacy loss from distributing HyperLogLog sketches.

We also are able to form some general conclusions about the expected privacy of HyperLogLog sketches. As mentioned above the prevalence ratio 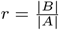, where *A* and *B* are respectively the background population and query population can be interpreted as the ratio of patients matching a query (e.g. ‘How many patients have been diagnosed with diabetes?’). Based on HLL, *m* is the number of buckets and *A_i_* and *B_i_* are the *i*th bucket in *A* and *B*. Figure 4 plots the number of buckets and prevalence rate against the estimated expected number of non ‘k-anonymized’ buckets and the number of buckets versus the percentage of the non ‘k-anonymous’ buckets. The two top plots are simply the number of non k-anonymous buckets against the number of buckets and varying the other parameters, but this turns out to not be the right set of variables to control.

**Fig. 4:**
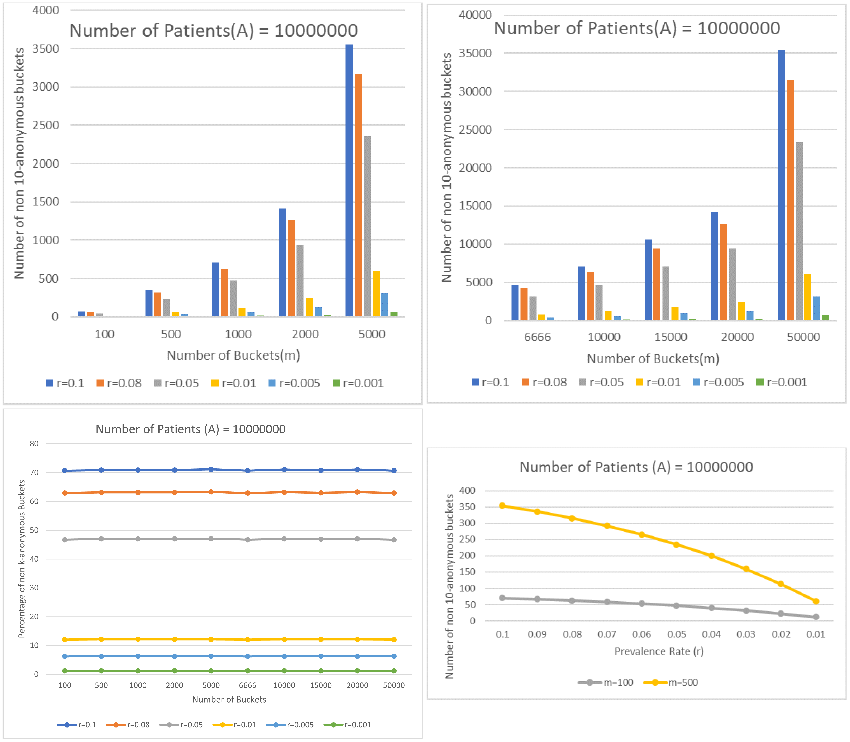
Expected number of non 10-anonymous buckets under different combinations of number of buckets (m) and prevalence rate (r) when total number of patients is 10^7^. (top) Number of non 10-anonymous buckets under different combinations of *m* (number of buckets) and *r* (prevalence rate) when total number of patients is 10^7^.(left bottom) However, the fraction of non 10-anonymous buckets remains constant as the number of buckets increase when the other variables are held fixed. (right bottom) It is the relationship to prevalence rate that is more complicated and nonlinear, as shown by focusing on the behavior for 100 and 500 buckets.

Instead, as evidenced by the lower left plot (Fig 4), a roughly constant fraction of the buckets are not k-anonymized when r is constant. This is unsurprising because as mentioned earlier, r is intuitively the number of background population members that could be used to hide each patient. Of course, random chance also plays a large role. More precisely, this constant is close to 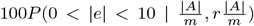 where 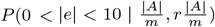 is the probability of that the number of collisions is greater than 0 and less than 10 when the bucket size is at the mean value 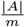. It is not quite equal for two reasons. The first reason is the obvious one, that we are using the approximations that form the subject of this manuscript. The second reason is that the single bucket size |*A*_1_| follows a Binomial distribution with mean 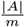 and 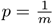. When |*A*| and 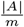 are big enough, we can get 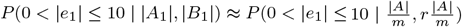 by concentrating |*A*_1_|, |*B*_1_| in a interval centered at the means, which is similar to what we did in Approximation A2, but simpler. However, when 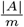 is not that big, for example, |*A*| = 100, *m* = 5, then 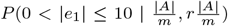 and *P*(0 < |*e*_1_| ≤ 10 | |*A*_1_|, *r*|*B*_1_|) will differ a lot for different value of |*A*_1_| and |*B*_1_|.

Now that it is clear that r is the value of primary importance, we see in the lower right plot of Fig 4 that the bigger the prevalence rate (*r*) is, the more buckets are non ‘k-anonymized’. This is because that bigger *r* means more overlap between set *A* and *B* and so is each pair of buckets *A_i_* and *B_i_*. Thus, the maximum number of zeros before the first one among all hash values represented in base-2 in *B_i_* is more likely equal to that in *A_i_*. Thus, a hospital IRB or clinical query system seeking to understand the 10-anonymity of a particular query can use a first-order approximation based only on *r*, without even needing to run our code. Indeed, they need only consult our lower-right plot in Fig 4 and scale to the size of their background population to determine that first-order approximation. This can be done without any code. When a more precise result is needed, however, our two Approximations can provide that answer in only a few minutes. Of course, if even that is insufficient, the practitioner may choose to directly measure the k-anonymity of a particular HLL sketch; this is not in the scope of this paper, but was empirically done in prior work (Yu and Weber, 2020).

## 6 Conclusion

In this article, we have developed a method to quickly compute the expected number of non ‘k-anonymous’ buckets in the HLL sketch. Because of the number of patients (denoted as |A| in our model) is too big to compute the precise expected value, we introduced two approximations based on concentration inequalities. In general, approximation A1 is suitable for the case when the expected value of single bucket size which is 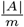 is ‘small’, for example, total number of patients(|*A*|) is 10^5^ and number of buckets (*m*) is 100 or total number of patients(|*A*|) is 10^7^ and number of buckets (*m*) is 10^5^. Approximation A2 is suitable for the case when 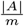 is ‘big’, for example, total number of patients(| A|) is 10^7^ and number of buckets (*m*) is 100 (see choice table in Result section).

By an appropriate choice of approximation method, we can control the computing time to under 300 seconds in almost all the cases. In other words, when an individual hospital is asked a query to return the aggregate counts based on sharing HLL sketches, we can compute the expected number of buckets which match fewer than 10 patients in the background population. If this number is too high, that is a signal to the clinical query system that the particular query is unsafe to release using HLL sketches. It is then up to the clinical query system to decide whether to fall back on another aggregation method, or if they should simply not respond to the query.

Our results further give some guidance into the parameter ranges in which HLL sketches are likely to be safe to release. HLL sketches are especially useful for rare diseases, where the prevalence ratio in the population is low. Note that this is in marked contrast to sending raw counts, where rare diseases are precisely the least k-anonymous. Thus, HLL sketches fill a complementary role.

Ultimately, our work is primarily useful in contexts where federated clinical query systems are used in biomedical research. The past year has seen increasing amounts of data centralization to combat the Covid-19 pandemic. The cost to privacy has been accepted because of the urgent clear and present need. However, in the future post-pandemic era as the pendulum swings the other direction, privacy may again take center stage. We hope that our work will be useful in analyzing the privacy consequences of distributed query systems and help inform policy-makers and institutional IRBs about the privacy-utility trade-offs at hand.

